# Methionine antagonizes *para*-aminosalicylic acid activity via affecting folate precursor biosynthesis pathway in *Mycobacterium tuberculosis*

**DOI:** 10.1101/319038

**Authors:** Michael D. Howe, Shannon L. Kordus, Malcolm S. Cole, Allison A. Bauman, Courtney C. Aldrich, Anthony D. Baughn, Yusuke Minato

## Abstract

*para*-Aminosalicylic acid (PAS) is a second-line anti-tubercular drug that is used for the treatment of drug-resistant tuberculosis (TB). PAS efficacy in the treatment of TB is limited by its lower potency against *Mycobacterium tuberculosis* relative to many other drugs in the TB treatment arsenal. It is known that intrinsic metabolites, such as *para*-aminobenzoic acid (PABA) and methionine, antagonize PAS and structurally related anti-folate drugs. While the basis for PABA-mediated antagonism of anti-folates is understood, the mechanism for methionine-based antagonism remains undefined. In the present study, we used both targeted and untargeted approaches to identify factors associated with methionine-mediated antagonism of PAS activity. We found that synthesis of folate precursors as well as a putative amino acid transporter play crucial roles in this process. We also discovered that intracellular biotin confers intrinsic PAS resistance in a methionine-independent manner. Collectively, our results demonstrate that methionine-mediated antagonism of anti-folate drugs occurs through sustained production of folate precursors.

## Introduction

*Mycobacterium tuberculosis* is responsible for approximately 10.4 million new cases of active tuberculosis (TB) and 1.3 million deaths annually (WHO, 2017). While TB chemotherapeutic intervention is highly successful in curing drug-susceptible TB infections, therapy is challenging, in part, because it requires a minimum of 6 months of treatment with drugs associated with adverse reactions. In addition, the emergence of drug-resistant strains of *M. tuberculosis* has dramatically increased the complexity and cost of TB treatment (Gandhi et al. 2006, Gehre et al. 2016). Therefore, the development of more efficacious TB chemotherapy regimens is imperative to improve treatment outcomes.

*para*-Aminosalicylic acid (PAS) was the second drug to be developed exclusively for TB chemotherapy (Lehmann 1946). Although PAS was a cornerstone agent of early multidrug TB therapies, the introduction of more potent anti-tubercular agents into TB treatment regimens greatly diminished its usage (Minato et al. 2015). Emergence of *M. tuberculosis* strains with resistance to first-line anti-tubercular agents led to the resurgence of PAS as an second-line drug to treat infections that failed to respond to standard short-course therapy (Donald and Diacon 2015). However, compared to many other anti-tubercular drugs, PAS is less potent and is associated with a high rate of gastrointestinal distress which limits its use to the treatment of multi-drug resistant TB for which there are few other treatment options (Zumla et al. 2013). Thus, it is important to develop novel strategies to enhance PAS potency, limit adverse reactions and improve treatment success rates.

Until recently, little was known regarding the mode of action of PAS. PAS is a selective antimetabolite of the *M. tuberculosis* folate metabolic pathway acting as a structural analog of the folate precursor *para*-aminobenzoic acid (PABA) (**Figure 1**) (Chakraborty et al. 2013, Minato, Thiede, Kordus, McKlveen, Turman and Baughn 2015). PAS is sequentially converted to 2’-hydroxy-7,8-dihydropteroate and 2’-hydroxy-7,8-dihydrofolate by enzymes in the *M. tuberculosis* folate metabolic pathway (**Figure 1**). 2’-hydroxy-7,8-dihydrofolate has been shown to potently inhibit *M. tuberculosis* dihydrofolate reductase (DHFR), the final step in synthesis of tetrahydrofolate (Dawadi et al. 2017, Minato, Thiede, Kordus, McKlveen, Turman and Baughn 2015, Zhao et al. 2014, Zheng et al. 2013). Since PAS and PABA are comparable substrates for the folate biosynthetic pathway, supplementation of *M. tuberculosis* cultures with PABA antagonizes the inhibitory activity of PAS by outcompeting for ligation to 6-pyrophosphomethyl-7,8-dihydropterin (DHPPP) by dihydropteroate synthase (DHPS) (Youmans et al. 1947). We previously reported that intracellular PABA mediates intrinsic resistance to PAS in *M. tuberculosis*, and disruption of this critical node in folate biosynthesis can potentiate antifolate action, including that of sulfa drugs (Thiede et al. 2016).

**Figure 1.**
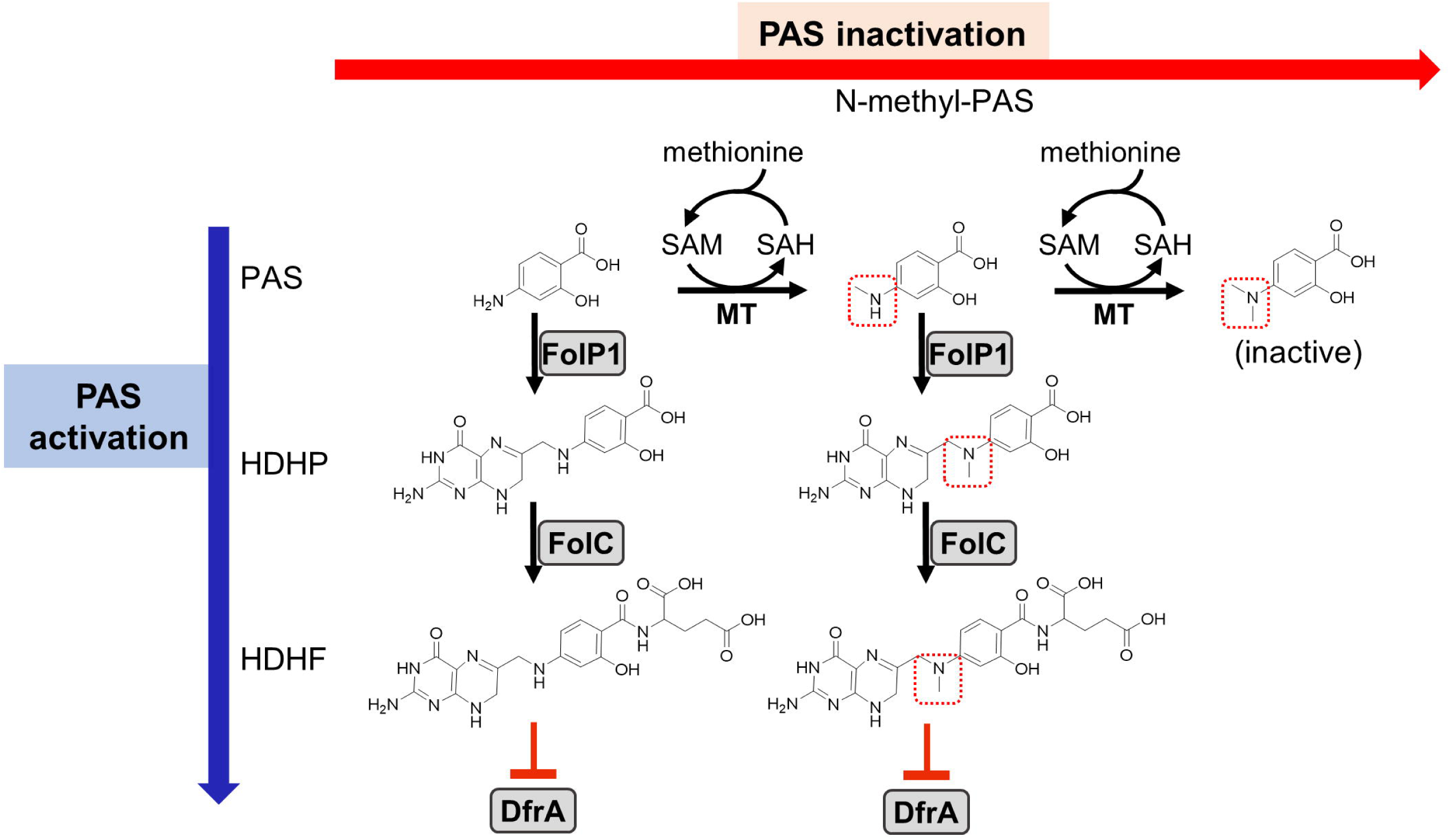
Previously proposed models for PAS activation and methionine-mediated PAS inactivation. As indicated on the left, PAS is a prodrug that must be activated by the *M. tuberculosis* folate biosynthetic pathway. PAS is incorporated in lieu of PABA by FolP1 and glutamoylated by FolC to form the antimetabolite HDHF which inhibits DfrA activity (indicated as red blunted arrows) (Dawadi, Kordus, Baughn and Aldrich 2017, Minato, Thiede, Kordus, McKlveen, Turman and Baughn 2015, Zhao, Wang, Erber, Luo, Guo, Yang, Gu, Turman, Gao, Li, Cui, Zhang, Bi, Baughn, Zhang and Deng 2014, Zheng, Rubin, Bifani, Mathys, Lim, Au, Jang, Nam, Dick, Walker, Pethe and Camacho 2013). Previous work has identified N-methyl and N,N-dimethyl PAS species (methyl groups indicated with dotted red boxes) in metabolite extracts from *M. tuberculosis* treated with PAS (Chakraborty, Gruber, Barry, Boshoff and Rhee 2013). As N,N-dimethylation of PAS prevents incorporation by FolP1, the resulting metabolite is inactive (shown on the right). This activity is presumed to be dependent upon an as of yet unidentified SAM-dependent methyltransferase(s). Supplementation with methionine may increase SAM pools, which could be utilized by the methyltransferase(s) to inactivate PAS, thus conferring resistance. Abbreviations: PAS, para-aminosalicylic acid; PABA, para-aminobenzoic acid; HDHP, 2’-hydroxy-7,8-dihydropteroate; HDHF, 2’-hydroxy-7,8-dihydrofolate; FolP1, dihydropteroate synthase; FolC, dihydrofolate synthase; DfrA, dihydrofolate reductase; MT, methyltransferase; SAM, S-adenosyl methionine; SAH, S-adenosyl homocysteine.

Methionine is a potent antagonist of PAS in *M. tuberculosis* (Hedgecock 1956), yet, the basis for this antagonism remains poorly understood. Because disruption of the folate pathway in *M. tuberculosis* results in depletion of metabolites within multiple essential folate-dependent pathways (Chakraborty, Gruber, Barry, Boshoff and Rhee 2013, Nixon et al. 2014), supplementation with methionine alone is not expected to recover loss of folate pathway integrity. A recent study showed that PAS can be converted to *N*-methyl and *N*,*N*-dimethyl PAS species within *M. tuberculosis* cells (**Figure 1**) (Chakraborty, Gruber, Barry, Boshoff and Rhee 2013). *N*-methyl-PAS retains activity against *M. tuberculosis*, while *N*,*N*-dimethyl-PAS shows no anti-tubercular activity since the resulting tertiary amine is incapable of nucleophilically reacting with DHPPP during the first step of PAS bioactivation (**Figure 1**). Since addition of methionine can potentially enhance the ability of *M. tuberculosis* to methylate PAS by increasing *S*-adenosylmethionine (SAM) abundance, it is possible that methionine promotes inactivation of PAS through *N*,*N*-dimethylation by an unidentified methyltransferase.

In the present study we screened approximately 10,000 independent *Mycobacterium bovis* BCG transposon insertion mutants (BCG::*himar1*) to identify genetic determinants associated with methionine-mediated PAS antagonism. In parallel to analysis of BCG::*himar1* mutants, we characterized factors that affect PAS susceptibility in *M. tuberculosis* for their involvement in methionine-mediated PAS antagonism. Our findings reveal the importance of folate precursor biosynthesis and methionine transport in methionine-mediated PAS antagonism.

## Materials and Methods

### Chemical Reagents

All chemical reagents except for 2’-hydroxy-pteroate (pterin-PAS) were purchased from Sigma-Aldrich. Pterin-PAS was synthesized by Drs. Richard Lee and Ying Zhao at St Jude Children’s Research Hospital by using a similar synthesis method reported elsewhere (Zhao et al. 2016).

### Bacterial Strains and Growth Conditions

Bacterial strains utilized in this study are described in **Table 1**. Unless otherwise indicated, Mycobacterial strains were grown in Middlebrook 7H9 liquid medium supplemented with tyloxapol (0.05% vol/vol) or on Middlebrook 7H10 agar plates. For *M. bovis* BCG and *M. tuberculosis* H37Ra, oleate-albumin-dextrose-catalase (OADC; Becton Dickinson 10% vol/vol) and glycerol (0.2% vol/vol) were supplemented to Middlebrook 7H9 and Middlebrook 7H10. For *Mycobacterium smegmatis* mc^2^155, Middlebrook 7H9 and Middlebrook 7H10 was amended with dextrose (0.2% vol/vol). *Escherichia coli* DH5αλ*pir* was grown in LB broth or on LB agar plate. When necessary, kanamycin or hygromycin were added to media at 50 µg/ml and 150 µg/ml respectively for selection of mycobacterial and *E. coli* strains.

**Table 1.**
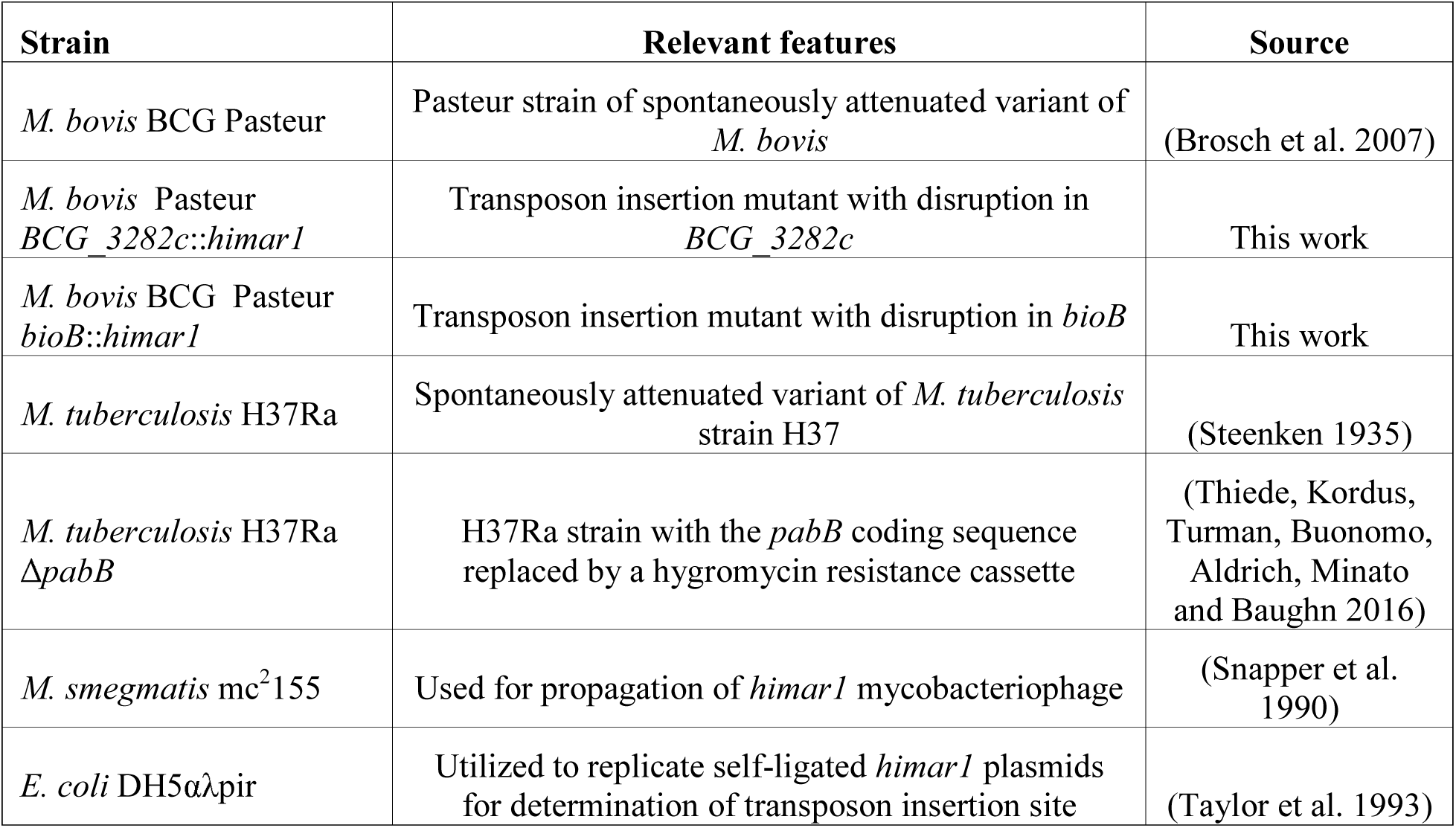
List of bacterial strains used in this study

For sulfur utilization studies, a modified sulfate-free Sautons medium (Allen 1998) was prepared with all inorganic sulfate salts (MgSO_4_ and ZnSO_4_) replaced with inorganic chloride salts (MgCl and ZnCl) keeping the concentrations of Mg^2+^ and Zn^2+^ ions the same. For the characterizations of the biotin auxotroph mutant, biotin-free 7H9 medium was prepared. The biotin-auxotrophic strain, *M. bovis* BCG *bioB*::*himar1*, was maintained in the biotin-free 7H9 medium supplemented with 0.5 µg/ml biotin. For the characterizations of the PABA auxotroph mutant, 7H9 medium was prepared in glassware that was baked at 300°C for one hour to remove residual PABA before use. The PABA-auxotrophic strain H37Ra Δ*pabB* was maintained in PABA-free 7H9 medium supplemented with PABA (10 ng/ml).

### Construction and Screening *M. bovis* BCG::*himar1* Mutant Library

The phAE180 mycobacteriophage containing a *mariner* transposable element, *himar1*, with a kanamycin resistance cassette was used to transduce *M. bovis* BCG creating a library of *BCG*::*himar1* mutants as described previously (Kriakov et al. 2003, Rubin et al. 1999). Transduced cells were plated onto 7H10 agar containing kanamycin and 10 µg/ml methionine. Approximately 10,000 mutant strains were screened by picking and patching onto 7H10 agar supplemented with methionine (Met plates) and onto 7H10 agar plates additionally amended with 5 µg/ml PAS (Met-PAS plates). Mutant strains that grew on the Met plates, but were inhibited for growth the Met-PAS plates, were selected for secondary screening following the same protocol. PAS susceptibility was assessed for strains that passed the secondary screen. *himar1* insertion sites were determined as previously described (Rubin, Akerley, Novik, Lampe, Husson and Mekalanos 1999). Briefly, extracted genomic DNA was digested with *Bss*HII and self-ligated to produce circular DNAs. The circularized DNAs that contained *ori6K* from a part of *himar1* transposon were used to transform *E. coli* DH5αλpir. Plasmids were purified from the transformants. Sequences of genomic DNA adjacent to the 3’ end of the *himar1* transposon insertion site were determined by Sanger sequencing (performed by Eurofins) using the KanSeq_Rev (5’-GCATCGCCTTCTATCGCCTTC-3’) primer (Baughn et al. 2010). Insertion site locations were determined by aligning the resulting sequence files with the *M. bovis* BCG Pasteur genome sequence (GenBank accession number NC_008796).

### Determination of Minimum Inhibitory Concentrations

The minimum inhibitory concentrations (MIC) of anti-tubercular compounds were determined as previously described (Dillon et al. 2014). Briefly, for determination of the MIC in liquid culture, 2-fold dilution series of drugs in 7H9 medium were prepared. Logarithmically growing Mycobacterium strains were inoculated into the drug-containing 7H9 medium in 30-ml square bottles (Nalgene) to an optical density (OD_600_) of 0.01. OD6_00_ were measured after shaking (100 rpm) at 37ºC for 14 days. The liquid MIC_90_ was defined as the minimum concentration of drug required to inhibit at least 90% of growth relative to growth in the no-drug control cultures. For determination of the agar plate MIC, logarithmically growing *M. bovis* BCG strains were serially-diluted and inoculated onto 7H10 agar plates containing drug in 2-fold dilution series. The agar plate MIC was determined by visually inspecting growth relative to growth on the no-drug control plates after grown at 37ºC for 21 days. All anti-tubercular compounds employed in this study were dissolved in DMSO. The highest concentration of DMSO in the growth media was 2.5%.

### Analysis of Growth Kinetics

Logarithmically growing Mycobacterium strains were washed twice in an equal volume of fresh medium. Cells were diluted to an OD_600_ of 0.01 in 30-ml square bottles (Nalgene) and supplements with or without drug were added at the described concentrations. Cultures were shaken (100 rpm) and OD_600_ were measured at various time points over a 14-day time-course.

### Methionine Utilization Experiments

*M. bovis* BCG strains were grown to mid-log phase in 7H9 broth and washed twice with sulfate-free Sautons medium. Resuspended cells were diluted to an OD_600_ of 0.01 in sulfate-free Sautons medium. Cultures were then incubated for 5 days to exhaust remaining sulfur. Exhausted cells were aliquoted into 30-ml square bottles (Nalgene) and sulfur-containing metabolites were added at the given concentrations. Cultures were incubated at 37ºC and shaken (100 rpm). The fold-change in OD_600_ (as a ratio of the final OD_600_/initial OD_600_) was assessed following 1 week of incubation after the addition of metabolites.

## Results

### Identification of *M. bovis* BCG genes involved in methionine-mediated antagonism of PAS

A library of *M. bovis* BCG transposon insertion mutant strains was constructed using the phAE180 mycobacteriophage containing a *mariner*-family transposable element. To identify genes associated with methionine-mediated PAS antagonism, approximately 10,000 BCG::*himar1* mutants were screened following the approach outlined in **Figure 2**. Determination of the PAS MIC on 7H10 agar plates confirmed that 0.25 µg/ml was sufficient to fully inhibit growth of the *M. bovis* BCG parental strain. Screening was then undertaken on 7H10 agar plates containing 10 µg/ml methionine and 5 µg/ml PAS (Met-PAS plate). Growth of *M. bovis* BCG on Met-PAS plates was identical to growth seen on control 7H10 plates, which confirmed methionine-mediated PAS antagonism.

**Figure 2.**
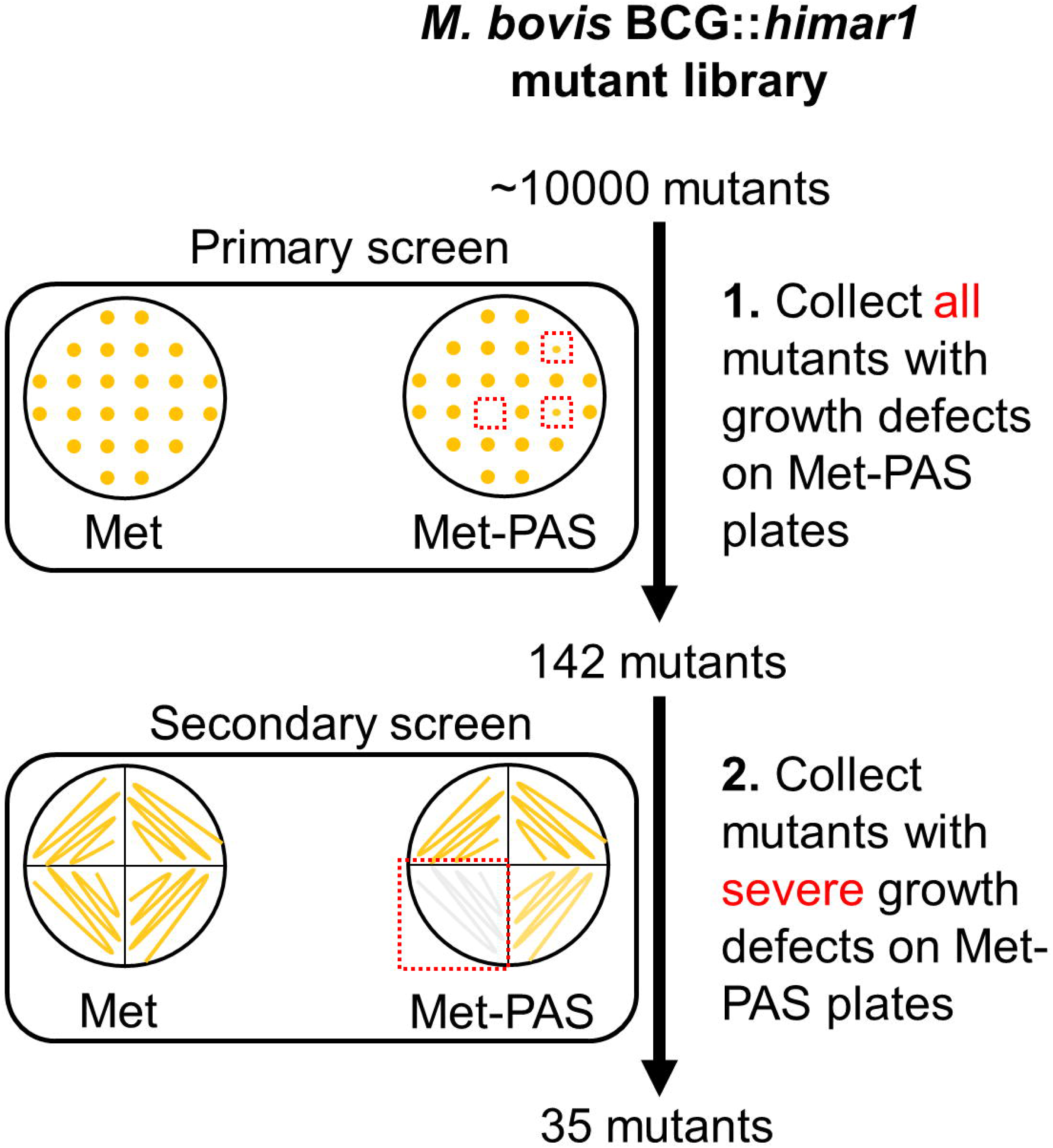
Schematic representation of genome-wide transposon mutagenesis of *M. bovis* BCG and Met-PAS screening method. *M. bovis* BCG::*himar1* mutants (approximately 10,000 mutants) were patched onto Met and Met-PAS plates. Colonies with observable growth defects on Met-PAS plates were subjected to secondary screening. These 35 mutants that passed the secondary screening were collected and insertion site locations were determined.

BCG::*himar1* insertion mutants which exhibited observable growth inhibition on the Met-PAS plates in comparison to the growth on 7H10 agar plates containing 10 µg/ml methionine (Met plate) were isolated. We then identified the *himar1* insertion sites within the 35 BCG::*himar1* mutants that had reproducible growth defects on Met-PAS plates compared to the growth on Met plates **(Figure 2 and Table 2)**. Among these mutants, one strain with a *himar1* insertion located within *BCG_3282c*, encoding a putative amino acid/polyamine/organocation (APC) superfamily transporter (Elbourne et al. 2017, Jack et al. 2000), showed the most severe growth defect on Met-PAS plates suggesting BCG_3282c plays a major role in methionine-mediated antagonism of PAS. We also assessed the susceptibility of each mutant strain to PAS by measuring PAS MICs on 7H10 agar plates **(Table 2)**. We observed the BCG_3282c mutant possessed wild-type PAS susceptibility suggesting this mutation is associated exclusively with methionine-mediated PAS antagonism.

**Table 2.**
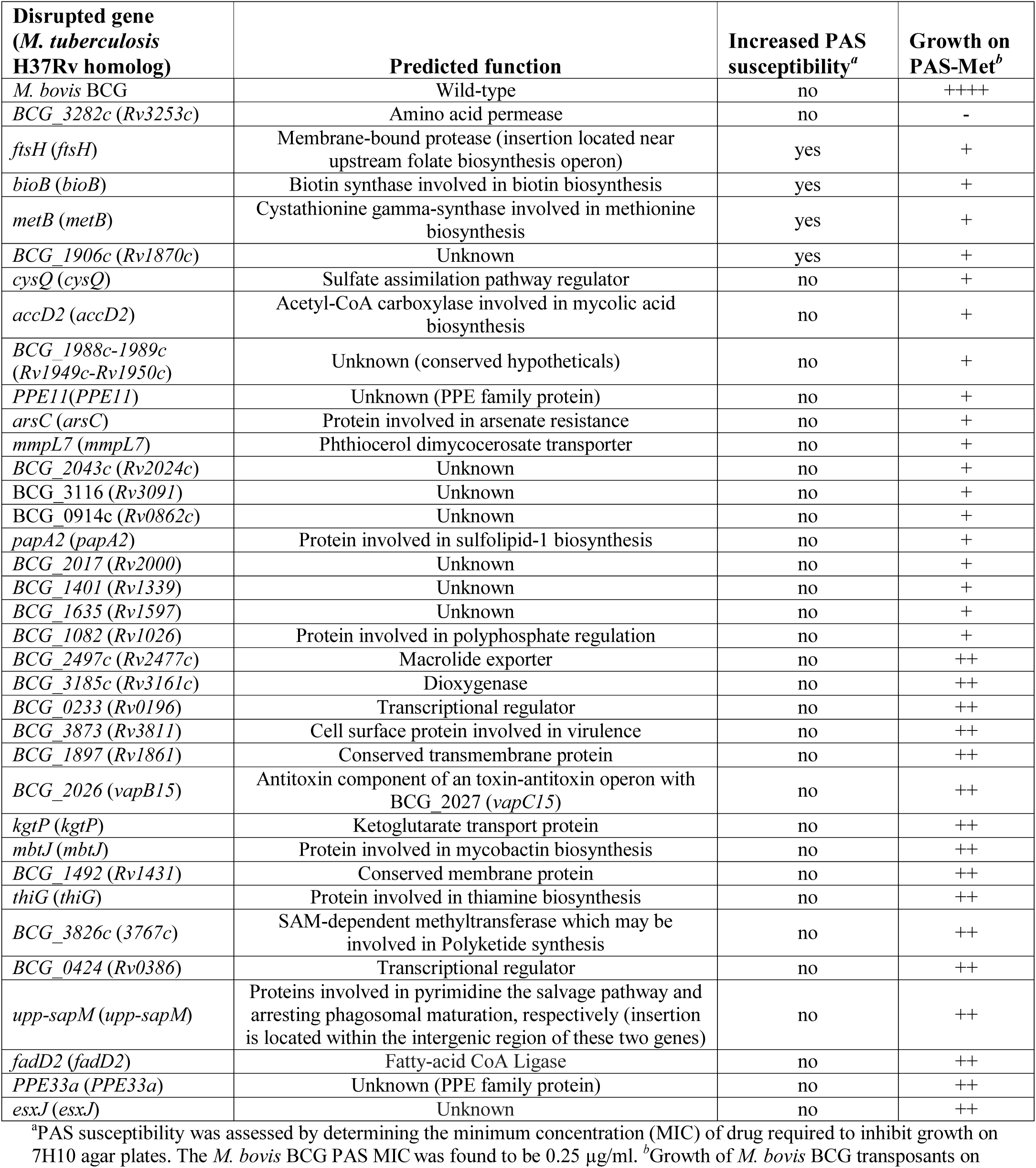
Sequence-validated gene insertions that affect PAS susceptibility in the presence or absence of antagonistic concentrations of methionine

Although most mutant strains that were analyzed showed a similar level of PAS tolerance as the parent *M. bovis* BCG, four mutants (with transposon insertions in *bioB*, *ftsH*, *metB*, and *BCG_1906c*) were found to be more susceptible to PAS in the absence of methionine, indicating that the disrupted genes may be involved in intrinsic resistance to PAS **(Table 2)**.

### BCG_3282c is essential for methionine-mediated antagonism of PAS in *M. bovis* BCG

Based upon the observation that methionine failed to antagonize PAS activity in the *BCG_3282c* mutant, we further characterized the function of this gene. The *himar1* insertion was located near the 5’ end of the coding region for *BCG_3282c* resulting in a 401 residue truncation of the 495 residue coding sequence, suggesting functional gene disruption by *himar1* insertion. Similar to the majority of transporters within the APC superfamily, BCG_3282c is predicted to possess 12 transmembrane α-helical spanners (Elbourne, Tetu, Hassan and Paulsen 2017). *BCG_3282c* is also highly conserved in the Mycobacterium genus, sharing 100% sequence identity with numerous *M. tuberculosis* complex organisms including *Rv3253c*, an ortholog from the standard virulent reference strain H37Rv. However, no close orthologs of BCG_3282c have been structurally or functionally characterized thus far.

To confirm whether BCG_3282c disruption altered methionine antagonism, PAS susceptibility testing was conducted in liquid medium **(Figure. 3A)**. Growth of both wild type *M. bovis* BCG and the *BCG_3282c*::*himar1* strain was severely inhibited by 5 µg/ml PAS. Addition of methionine restored growth during PAS treatment of wild-type *M. bovis* BCG in a dose-dependent manner. In contrast, growth of the *BCG_*3282c::*himar1* strain was inhibited by PAS even in the presence of 10 µg/ml methionine. PABA, another PAS antagonist, reversed PAS-mediated growth inhibition in both the wild type *M. bovis* BCG and the *BCG_3282c*::*himar1* strain, validating that the *BCG_3282c*::*himar1* strain is specifically impaired for methionine-mediated PAS antagonism.

**Figure 3.**
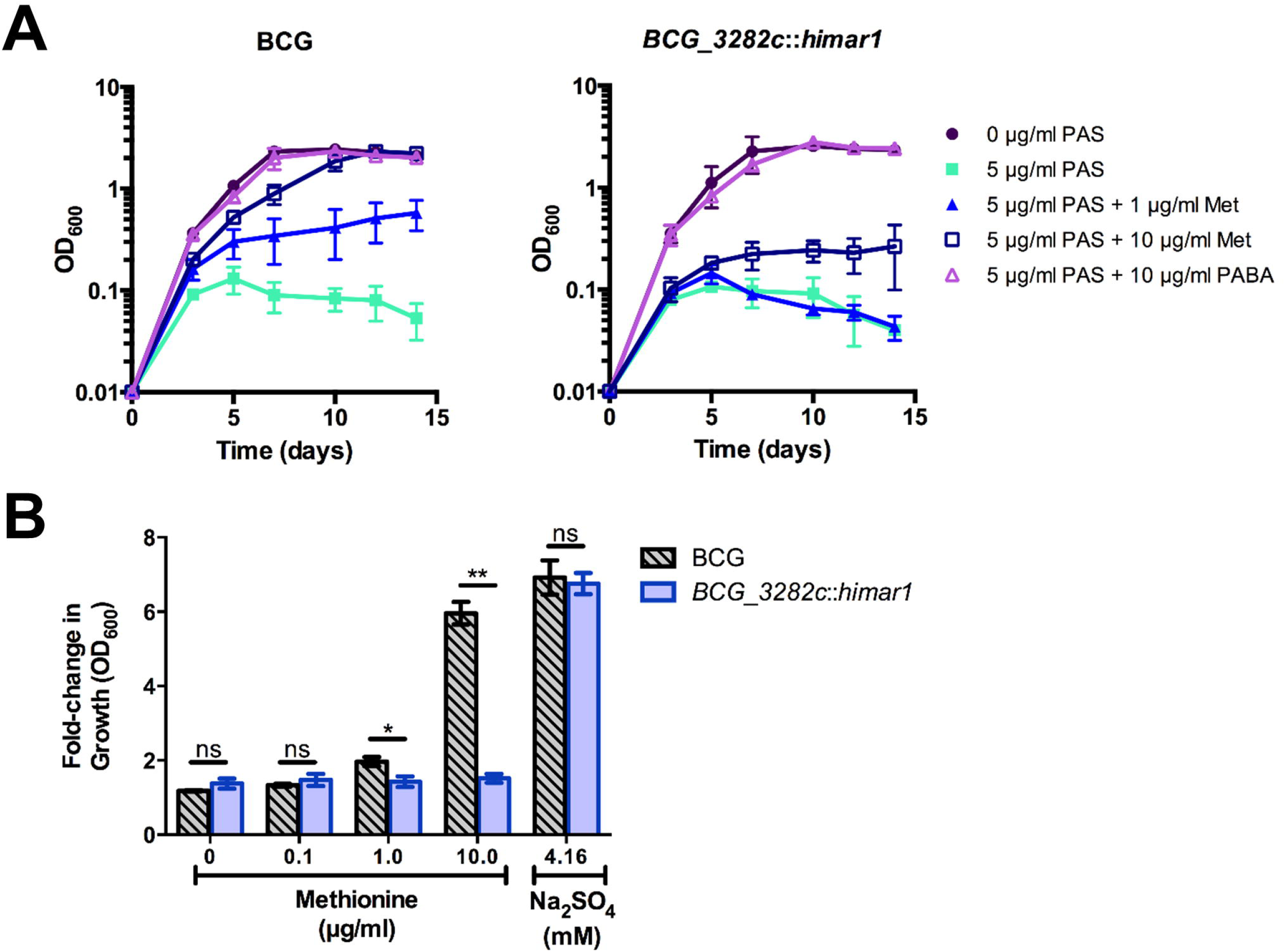
BCG_3282c is essential for methionine-mediated PAS antagonism and utilization of sulfur derived from methionine in *M. bovis* BCG. **(A)** Growth kinetics of *M. bovis* BCG *BCG_3282c*::*himar1* and *M. bovis* BCG wild type during PAS exposure when antagonistic metabolites are added. Growth was assessed by OD_600_ readings every 2-3 days. **(B)** *M. bovis* BCG strains were grown to an OD_600_ of approximately 0.5, washed three times to remove residual sulfate with sulfate-free Sautons medium, and resuspended in sulfate-free Sautons medium to a starting OD_600_ of 0.01 and cells were starved for sulfur for 5 days. Following the exhaust period, sulfur-sources were added, and cells were incubated for 7 days to resume growth. The fold-change in growth was assessed as a ratio of the final OD_600_ over the starting OD_600_ following the exhaust period (final OD_600_/starting OD_600_). p-values of pairwise comparisons (denoted by lines) were calculated using the Student t test. *, p < 0.05, **, p.<.0.005, ns indicates no significant difference (p > 0.05). **(A,B)** Error bars denote standard deviation and are representative of 3 separate experiments.

Because methionine antagonism is selectively perturbed in the BCG_3282c::*himar1* mutant, we hypothesized that BCG_3282c may be the methionine transporter. *M. tuberculosis* is known to utilize reverse transsulfuration to assimilate sulfur from methionine which can serve as the sole source of sulfur for this bacterium (Wheeler et al. 2005). Therefore, we tested whether disruption of *BCG_3282c* would affect the ability of the bacilli to assimilate sulfur derived from methionine. When *M. bovis* BCG and the *BCG_3282c*::*himar1* disruption strain were grown in sulfate-free Sautons medium, growth of both strains was limited (maximum OD_600_ = 0.3). Upon addition of sodium sulfate to the medium, both strains resumed growth and achieved typical growth yields confirming these strains were previously starved for sulfur **(Figure. 3B)**. When methionine was added to sulfur starved *M. bovis* BCG, growth also resumed in a dose-dependent manner producing similar growth yields as compared to the addition of sulfate alone. In contrast, growth of the *BCG_3282c*::*himar1* disruption strain could not be restored in the presence of methionine as the sole source of sulfur, indicating that methionine is a transport substrate of BCG_3282c. These findings indicate antagonism requires methionine import across the cell membrane via BCG_3282c.

### PABA biosynthesis is indispensable for methionine-mediated PAS antagonism in *M. tuberculosis*

Our large-scale screening failed to identify genes directly involved in methionine-mediated PAS antagonism. Thus, it is possible that genes involved in this process are redundant or are essential for *M. bovis* BCG survival *in vitro*. Because addition of methionine can increase SAM levels and many *M. tuberculosis* SAM-dependent methyltransferase genes are essential, we investigated whether the ability to methylate PAS plays a role in methionine-mediated PAS antagonism. To test this, we evaluated whether methionine can antagonize the activated PAS species, 2’-hydroxy-pteroate (pterin-PAS), in *M. bovis* BCG. It is known that *N*,*N*-dimethyl-PAS has no anti-tubercular activity, presumably because *N*,*N*-dimethyl-PAS cannot react with DHPPP during the first step of PAS bioactivation (**Figure 1**). Thus, once PAS is activated to pterin-PAS, *N-*methylation should not affect its anti-tubercular activity. We confirmed pterin-PAS was active against wild-type *M. bovis* BCG at a comparable molar concentration to PAS (**Table 3**). Surprisingly, pterin-PAS was still potently antagonized by methionine suggesting methionine-mediated PAS antagonism does not occur by methylation of PAS to inhibit PAS bioactivation.

**Table 3.** Antagonism of PAS and pterin-PAS by methionine

| Strain | PAS MIC<sub>90</sub><sup>a</sup> |  | Pterin-PAS MIC<sub>90</sub> |  |
| --- | --- | --- | --- | --- |
|  | - Met | + Met<sup>b</sup> | - Met | + Met |
| *M. bovis* BCG | 1 (6.53) | >250 (1630) | 5 (15.1) | >20 (61.5) |
| *M. tuberculosis* H37Ra | 0.6 (3.92) | >250 (1630) | ND | ND |
| *M. tuberculosis* H37Ra ΔpabB | 0.15 (0.98) | 0.3 (1.96) | ND | ND |

<sup>a</sup>MIC<sub>90</sub> is defined as the minimum concentration of drug required to restrict at least 90% of growth relative to growth seen in the no-drug control cultures. MIC<sub>90</sub> are shown in µg/ml (µM).
ND, not determined; Met, methionine; PAS, *para*-aminosalicylic acid.
<sup>b</sup>Methionine was supplemented at 10 µg/ml.

It is also known that intracellular PABA levels affect PAS susceptibility in *M. tuberculosis* (Thiede, Kordus, Turman, Buonomo, Aldrich, Minato and Baughn 2016). Since PABA biosynthesis is essential for Mycobacterium survival *in vitro*, we hypothesized that methionine may affect PAS activity. PabB, aminodeoxychorismate synthase, is one of the essential enzymes required to convert chorismate to PABA in *M. tuberculosis* (**Figure 4A**). Consequently, a *M. tuberculosis* H37Ra *pabB* deletion strain is a PABA auxotroph and relies upon exogenous sources of PABA for growth (**Figure 4B**). The folate precursor dihydropteroate is produced from PABA and DHPPP (**Figure 4A**). We found that pteroic acid, an oxidized form of dihydropteroate can also support the growth of the *M. tuberculosis* H37Ra *pabB* deletion strain (**Figure 4B**). As expected, unlike PABA and pteroic acid, methionine did not support the growth of the *M. tuberculosis* H37Ra *pabB* deletion strain indicating that methionine alone is insufficient to fulfill cellular folate requirements in PABA starved *M. tuberculosis* cells. Using the *M. tuberculosis* H37Ra Δ*pabB* deletion strain, we tested the requirement of PABA biosynthesis on methionine-mediated PAS antagonism. We observed that methionine potently antagonized PAS susceptibility in wild type *M. tuberculosis* H37Ra. In contrast, PAS susceptibility of the *M. tuberculosis* H37Ra Δ*pabB* deletion strain was not antagonized by the addition of methionine **(Table 3)**. Taken together, these data demonstrated that a functional PABA biosynthetic pathway is essential for methionine to antagonize PAS in *M. tuberculosis*.

**Figure 4.**
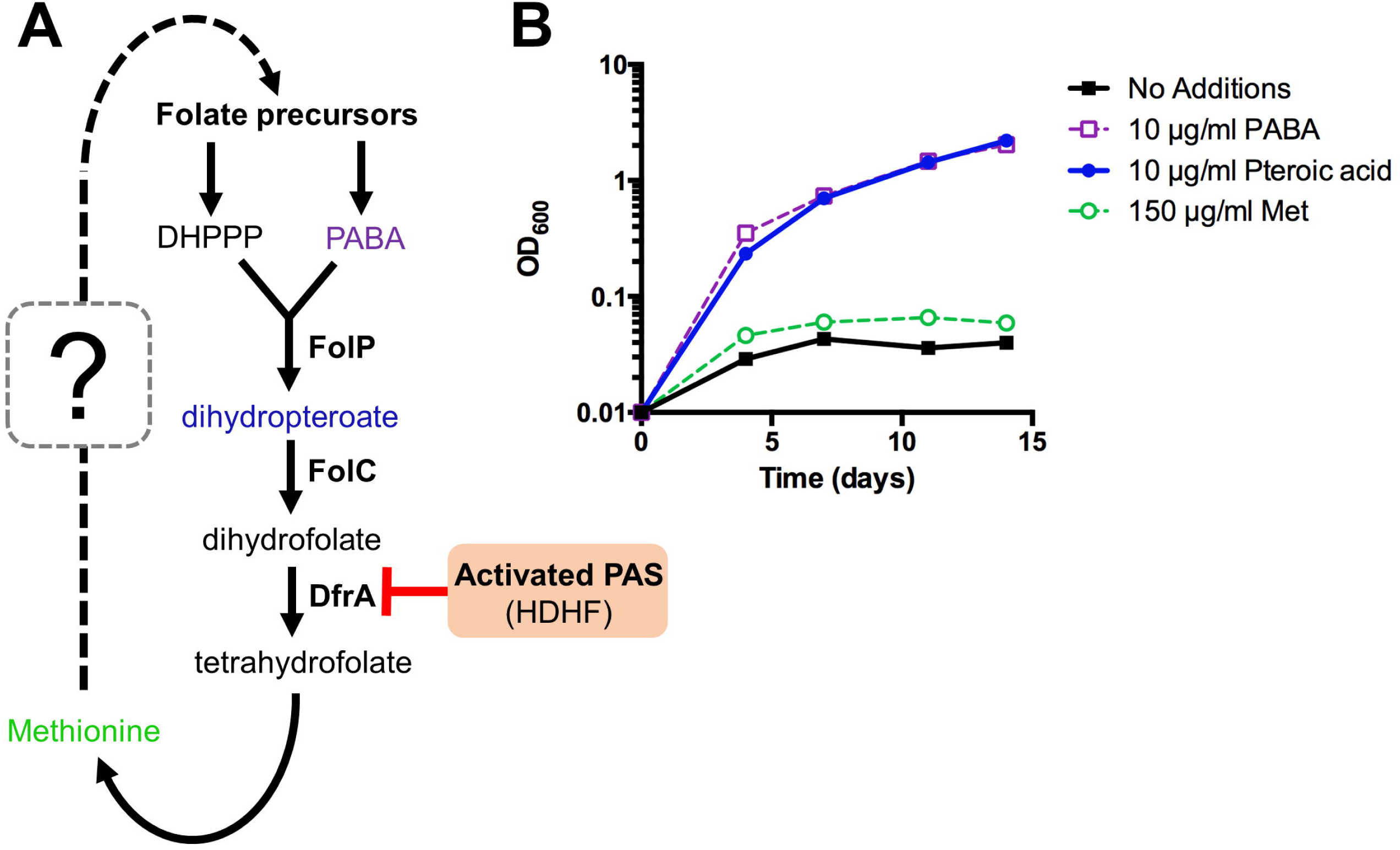
Methionine can affect but not bypass essentiality of upstream folate biosynthetic pathways in *M. tuberculosis*. **(A)** New working model of methionine-mediated PAS antagonism. **(B)** *M. tuberculosis ΔpabB* was grown to an OD_600_ of approximately 0.5, washed three times to remove residual PABA with PABA-free 7H9 medium, and resuspended in PABA-free 7H9 medium to a starting OD_600_ of 0.01 (3 × 10^6^ cells/mL). Cultures were then supplemented with the indicated metabolites and incubated for 14 days with OD_600_ readings taken at the given time points.

### Biotin cofactor biosynthesis is essential for intrinsic resistance to PAS and other anti-tubercular drugs

Our screening also identified several mutations that conferred increased susceptibility to PAS even in the absence of methionine. One strain, harboring a *himar1* insertion within *bioB*, encoding biotin synthase, showed increased susceptibility to PAS both in the presence and absence of methionine **(Table 2)**. BioB is a radical SAM-dependent enzyme required for the final step in the synthesis of biotin. We confirmed the *bioB*::*himar1* strain exhibited biotin auxotrophy, which could be chemically complemented by a minimum of 0.05 µg/ml biotin supplementation for restoration of growth **(Figure 5A)**. We speculated that susceptibility of the *bioB*::*himar1* strain to PAS was dependent upon intracellular concentrations of biotin. Thus, we examined the PAS susceptibility of the *bioB*::*himar1* strain using media containing minimal (0.05 µg/ml) or excess (5 µg/ml) concentrations of biotin **(Figure 5B)**. We observed the *bioB*::*himar1* strain was far more susceptible to PAS (8-fold decrease in MIC90) in minimal biotin medium, and that excess biotin medium was sufficient to restore susceptibility back to near wild-type levels. Interestingly, the *bioB*::*himar1* strain was also more susceptible to sulfamethoxazole (SMX) and rifampicin (RIF), but maintained wild-type susceptibility to isoniazid, indicating that alterations in susceptibility profiles are drug-specific **(Figure 5B)**.

**Figure 5.**
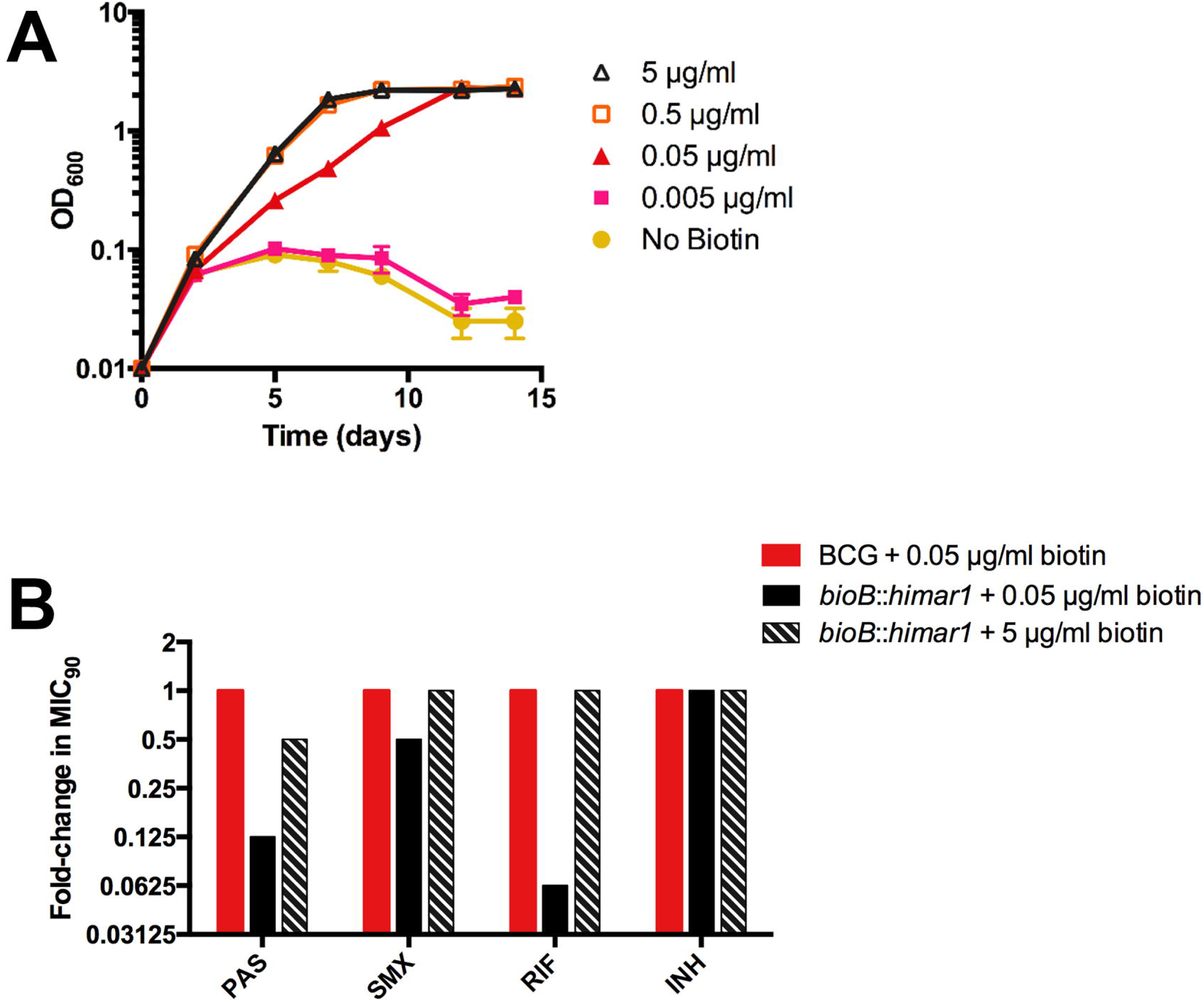
Disruption of *bioB* is growth inhibitory and potentiates drug action. **(A,B)** *M. bovis* BCG *bioB*::*himar1* was grown to an OD_600_ of approximately 0.5, washed three times to remove residual biotin with biotin-free 7H9 medium, and resuspended in biotin-free 7H9 medium to a starting OD_600_ of 0.01. Cultures were then supplemented with biotin and incubated for 14 days with OD_600_ readings taken at the given time points. Error bars denote standard deviation and are representative of 2 separate experiments. **(B)** *M. bovis* BCG and *bioB*::*himar1* were grown to an OD_600_ of approximately 0.5, washed three times to remove residual biotin, and resuspended in biotin-free 7H9 medium to a starting OD_600_ of 0.01. Cultures were then supplemented with biotin (0.05 and 5) and incubated for 14 days with OD_600_ readings taken at the given time points. MIC_90_ is defined as the minimum concentration of inhibitor required to restrict at least 90% of growth relative to growth seen in the no-drug control cultures. Abbreviations: PAS, *para*-aminosalicylic acid; SMX, sulfamethoxazole; RIF, rifampin; INH, isoniazid. Results shown are representative of 3 separate experiments.

## Discussion

Methionine is the only folate-dependent metabolite known to antagonize certain anti-folate drugs in *M. tuberculosis* and other bacterial species. Interestingly, anti-folate drugs antagonized by methionine are also antagonized by PABA, a folate precursor. Although the molecular mechanism of PABA-mediated anti-folate antagonism is well understood, how methionine antagonizes anti-folate drugs has yet to be elucidated. Our findings revealed that methionine-mediated PAS antagonism is linked to synthesis of folate precursors.

One strain isolated from our screen harboring a *himar1* disruption within the predicted amino acid permease *BCG_3282c* fully sensitized *M. bovis* BCG to PAS in the presence of normally antagonistic concentrations of methionine. In addition, the *himar1* disruption within *BCG_3282c* prevented *M. bovis* BCG from assimilating sulfur derived from methionine. BCG_3282c belongs to the APC superfamily of transporters and our data suggested that BCG_3282c is likely responsible for uptake of methionine *in vitro*. The most well-studied methionine transport system in bacteria is the MetD ABC transporter system of the methionine uptake transporter family found in numerous organisms including *E. coli* and even the closely related non-tubercular Mycobacterium, *Mycobacterium abscessus* (Gál et al. 2002). In *E. coli*, the MetD ABC transporter is encoded by the *metNIQ* gene cluster (Merlin et al. 2002). The *M. tuberculosis* complex has no known orthologs of this system, despite the known bioavailability of methionine in human and mouse serum (Lewis et al. 1980, Rivera et al. 1987). To our knowledge, this study represents the first characterization of a methionine transporter in the *M. tuberculosis* complex. Orthologues of BCG_3282c with high amino acid sequence similarities are found from *Gordonia sputi*, *Bacillus subtilis* and *Lactococcus lactis* and an orthologue from *L. lactis* has been shown to transport branched-chain amino acids, along with methionine (den Hengst et al. 2006). Existence of a conserved methionine transporter within the mycobacterium complex would be intriguing given that methionine/SAM biosynthesis is indispensable for survival of *M. tuberculosis* in murine and macrophage models of infection (Berney et al. 2015).

We also found that methionine-mediated PAS antagonism does not appear to occur through *N*,*N*-dimethylation by SAM-dependent methyltransferase(s). We addressed this possibility because *N*,*N*-dimethyl PAS, an inactive metabolite of PAS, was previously identified in metabolite extracts from PAS treated *M. tuberculosis* (Chakraborty, Gruber, Barry, Boshoff and Rhee 2013). In addition, a SAM-dependent methyltransferase (*Rv0560c*) is induced by salicylate and salicylate analogs, including PAS (Schuessler and Parish 2012). However, a recent report described that an unmarked in-frame deletion of *Rv0560c* in *M. tuberculosis* conferred no alteration in susceptibility to PAS, or other antimicrobials *in vitro* (Kokoczka et al. 2017). Consistent with this finding, our screen did not identify *Rv0560c*::*himar1* mutants. Together with our observation that pterin-PAS is also antagonized by methionine, we concluded that methionine-mediated PAS antagonism is not likely via PAS inactivation by *N*,*N*-dimethylation.

Importantly, methionine was unable to antagonize PAS in a *pabB* deletion mutant strain indicating that methionine-mediated PAS antagonism is dependent upon a functional PABA biosynthesis pathway. This finding is consistent with past and recent reports that methionine only antagonizes the anti-folate drugs that are also antagonized by PABA (Huang et al. 1997, Nixon, Saionz, Koo, Szymonifka, Jung, Roberts, Nandakumar, Kumar, Liao, Rustad, Sacchettini, Rhee, Freundlich and Sherman 2014, Zhao, Shadrick, Wallace, Wu, Griffith, Qi, Yun, White and Lee 2016, Zheng, Rubin, Bifani, Mathys, Lim, Au, Jang, Nam, Dick, Walker, Pethe and Camacho 2013). While the metabolic connections linking methionine to folate precursor biosynthesis remain to be determined, the DHPPP pathway has been shown to modulate susceptibility of *E. coli*, *Salmonella enterica* and *Burkholderia pseudomallei* to sulfamethoxazole (Li et al. 2017, Podnecky et al. 2017, Minato et al. 2018), which is predicted to be metabolically linked with methionine-mediated antagonism (Minato and Baughn 2017). Further, we recently demonstrated that the biosynthetic pathway to DHPPP is essential for methionine-mediated antagonism of sulfonamide action in *E. coli* (Minato et al. 2018).

One PAS-sensitive mutant strain with a disruption in biotin synthase (*bioB*) was found to be auxotrophic for the cofactor biotin. Characterization of this mutant confirmed that disruption of biotin biosynthesis could enhance susceptibility to PAS and rifampicin in biotin-limited conditions. In *M. tuberculosis*, biotin is a cofactor required for acyl-CoA-carboxylase (ACC) enzymes participating in key metabolic processes in lipid biosynthesis (Gago et al. 2011, Salaemae et al. 2011, Takayama et al. 2005, Woong Park et al. 2011). Biotin biosynthesis and protein biotinylation process have been targeted for novel drug development (Duckworth et al. 2011, Shi et al. 2013, Tiwari et al. 2018). On the basis of our *in vitro* findings, targeting biotin synthesis may promote accumulation of antimycobacterial drugs by disrupting cell envelope integrity, which could revitalize drug therapies that are unable to overcome the relatively impermeable cell envelope of *M. tuberculosis* at clinically relevant dosages. Indeed, it was recently reported that disruption of protein biotinylation potentiates rifampicin activity against *M. tuberculosis* (Tiwari, Park, Essawy, Dawadi, Mason, Nandakumar, Zimmerman, Mina, Ho, Engelhart, Ioerger, Sacchettini, Rhee, Ehrt, Aldrich, Dartois and Schnappinger 2018). It was previously reported that biotin has a vital role in methionine-mediated, PAS antagonism, such that supplementation with exogenous biotin was required to observe antagonism by methionine (Hedgecock 1956). However, our study found that biotin supplementation was non-essential for methionine to antagonize PAS in *M. bovis* BCG suggesting that the effect of biotin on PAS susceptibility is independent of the precise mechanism of antagonism, and the initial observations in *M. tuberculosis* by Hedgecock may be a strain specific phenotype.

In summary, the mechanistic basis of methionine-mediated PAS antagonism was examined. Over 30 novel modulators of PAS susceptibility were identified by *Himar1* transposon mutagenesis. However, with exception of the putative amino acid permease BCG_3282c, none of the functions identified were found to be directly involved in antagonism. Upon closer examination, *de novo* biosynthesis of PABA was determined as essential for methionine-mediated antagonism, revealing a previously unappreciated relationship between methionine and folate precursor synthesis. Further studies are needed to reveal the precise mechanism of this process. The results presented here also identified tractable drug targets within *M. tuberculosis* that could be exploited to enhance antimycobacterial drug action.

## Author Contributions

MDH, SLK, MSC, and AAB performed experiments. ADB, CCA and YM conceived the work. MDH, ADB and YM wrote the manuscript. All authors contributed to analyzing data and editing of the manuscript.

## Funding

This work was supported by a grant from the University of Minnesota Academic Health Center Faculty Research Development Program to ADB and CCA and by startup funds from the University of Minnesota to ADB.

## Conflict of Interest Statement

The authors declare that the research was conducted in the absence of any commercial or financial relationships that could be construed as a potential conflict of interest.

## Acknowledgements

We thank Nicholas D. Peterson for assistance in construction of a library of *M. bovis* BCG transposon insertion mutants. We thank Drs. Richard Lee and Ying Zhao of St Jude Children’s Research Hospital for the providing pterin-PAS.

